# Following embryonic stem cells, their differentiated progeny, and cell-state changes during iPS reprogramming by Raman spectroscopy

**DOI:** 10.1101/2020.05.02.074476

**Authors:** Arno Germond, Yulia Panina, Mikio Shiga, Hirohiko Niioka, Tomonobu M. Watanabe

## Abstract

To monitor cell state transition in pluripotent cells is invaluable for application and basic research. In this study, we demonstrate the pertinence of use non-invasive, label-free Raman spectroscopy to monitor and characterize the cell state transition of mouse stem cells undergoing reprogramming. Using an isogenic cell line of mouse stem cells, reprogramming from neuronal cells was performed, and we showcase a comparative analysis of single cell spectral data of the original stem cells, their neuronal progenitors, and reprogrammed cells. Neural network, regression models, and ratiometric analysis were used to discriminate the cell states and extract several important biomarkers specific to differentiation or reprogramming. Our results indicated that the Raman spectrum allowed to build a low dimensional space allowing to monitor and characterize the dynamics of cell state transition at a single cell level, scattered in heterogeneous populations. Ability of monitoring pluripotency by Raman spectroscopy, and distinguish differences between ES and reprogrammed cells is also discussed.

## Introduction

Pluripotent stem cells are defined as cells that are able to differentiate into all three embryonic layers and they hold enormous potential for future medical applications and basic research ^1, 2,3, 4^. Pluripotent stem cells can be directly obtained from a developing embryo, and in that case are called embryonic stem cells (ESCs), or can be artificially produced by somatic cell reprogramming and are then called induced pluripotent stem cells (iPSCs) ^5^. Reprogramming is a lengthy process that involves profound changes to the cells’ transcriptional, proteomic, metabolomic, etc. ^6^. Monitoring of the cell dynamics during reprogramming and recording the occurring changes is of utmost importance for both applied medicine and basic research as it can reveal important qualities of the process itself and of the cells going through the process. Conventional monitoring of the cell dynamics during reprogramming includes fluorescence microscopy/immunostaining for common pluripotency factors such as Nanog and Oct4, western blotting, and gene expression measurements by quantitative PCR or RNA sequencing. These approaches are informative in regard to pluripotency status, but they are destructive, cell-invasive and entail significant costs for evaluation of the cell quality.

A promising alternative to the above methods is Raman spectroscopy which is a non-invasive, low-cost and rapid single-cell imaging technique ^7, 8, 9^. Raman spectroscopy is a type of nonlinear vibrational spectroscopy whose output is a spectrum of molecules (molecular bounds) present within the sample being studied. Usually this “chemical fingerprint” is a complex mixture of convoluted signals from various molecules. Previous works have demonstrated that, however, that the major constituents of cells such as cytochromes, proteins, aromatic compounds, nucleic acids and lipids can be identified by this technique. Raman spectroscopy is increasingly used for the identification of cell types or dynamics across many biological systems ^10^, thanks to the rich signature obtained from cells.

The use of Raman spectroscopy in the stem cell biology has been investigated by several groups. Notingher and colleagues ^11^ first used Raman spectroscopy to examine live murine ESCs over the course of 16 days of differentiation. Chan and colleagues ^12^ used Raman spectroscopy to examine live human ESCs during differentiation into cardiomyocytes. They reported the changes in intensities of the RNA peak (~ 811 cm^-1^) and DNA peaks (~ 785, ~ 1090 cm^-1^) during differentiation, confirming the observation by Notingher and colleagues. Schulze *et al.* ^13^ analyzed hESCs obtained from dried colonies to characterize the differentiation process between ES cells and spontaneously differentiated cells. Using the spectral range of 690-1210 cm^-1^, authors revealed spectral differences and calculated a number of ratios of possible interest, such as a ratio between protein-related bands (~ 752cm^-1^) to nucleic acid bands (~ 786 cm^-1^) which may reflect the differentiation process. Our group previously showcased the large spectral information contained in the 700-1800 cm^-1^ fingerprint region, which was useful to visualize the transition of the cellular dynamics through multiple “states” of mouse differentiation ^10^.

These previous studies demonstrated that Raman spectroscopy is useful to monitor the process of differentiation in stem cells. From a system-biology point of view, the possibility to define a low dimensional space from spectral characteristics is particularly useful to monitor the dynamics of the differentiation process. In this space, different clusters of cells would represent different cellular states which have different characteristics. However, as far as we know, no study was performed on the reprogramming process as of today. A comparison between both differentiated and reprogrammed cells from the same isogenic cell line is therefore of interest to reveal the characteristics of spectral signatures of cells undergoing reprogramming by comparison to their differentiated states or by comparison to the embryonic cell state. In addition, it worth comparing if the ES and reprogrammed cells are, based on their spectral characteristic, associated to a similar state or not.

In this paper, we aim to assess the spectral differences and similarities of embryonic stem cells (ES), the differentiated progeny of these embryonic cells (called N31), and cells collected at three time points during reprogramming of N31 (hereafter, called N31d5, N31d10, N31d20). For this purpose, we carried out iPS reprogramming of mouse neural progenitors and obtained their Raman signature throughout the process. We characterized the pluripotency status of mouse cells during the iPS reprogramming using conventional methods such as immunostaining. Label-free spectroscopy and spectral analyses were then performed on living cells with the aim of finding spectral differences specific to each state. Machine learning analyses were used to classify cells with excellent accuracy and retrieve the cell state specific spectral information to characterize each cell state. Finally, ratiometric analysis revealed pairs of spectral bands representative of the cell states dynamics.

## Materials and Methods

### Cell culture

#### Embryonic stem cells

EB5 embryonic stem cells were purchased from RIKEN Cell Bank (Cell ID #AES0151) and kept in DMEM (Gibco, #11960) with the addition of 10% FBS (Gibco, #16141), 1% GlutaMAX-I (Gibco #35050), 1% NEAA (Gibco #11140), 1% Nucleosides (Millipore ES-008-D), 1% Sodium pyruvate (Sigma-Aldrich #S8636), 0.1% 2ME (Sigma-Aldrich #M7522), 1% Penicillin-Streptomycin (Sigma-Aldrich #P4333-100ML) and 0.1% LIF (Sigma-Aldrich #L5283). The EB5 cells were used for Raman imaging after two passages. *Differentiated cells*. N31 cell is a neural lineage derived from EB5 as described in Hikichi *et. al.* ^14^. The N31 clone was kindly provided by Prof. Shinji Masui from Kyoto University. To obtain N31, a doxycycline-inducible iPS OKSM system based on PiggyBac transposition was introduced into the EB5 cell population in order to facilitate iPS induction experiments ^14^. EB5 cells were differentiated for 7 days then seeded onto gelatin-coated T75 flask in 10 mL of NPC as single cells so as to obtain clones (for details, see ^14^). The resulting cell state, designated as clone N31, was maintained on gelatin-coated dishes in RHB-Basal medium (Clontech Takara, #Y40000) with NDiff^®^ N2 neural cell supplement (Clontech Takara, # Y40100) and 1% Penicillin-Streptomycin (Sigma-Aldrich #P4333-100ML). The N31 line was subsequently used for Raman imaging, representing a differentiated state.

#### N31 reprogramming

For converting the differentiated cells (N31 line) into iPS cells, we induced the iPS reprogramming process in the N31 line for the total duration of 15 days. On the day of reprogramming induction the RHB-Basal medium was replaced with serum-free, chemically defined iPS induction medium PSGro^®^ (StemRD, # PGro-500) with the addition of 10 ug/ml doxycycline, and the medium was exchanged every day until the reprogramming was complete. When ready, cell colonies were harvested, dissociated, and single cells were directly used for Raman imaging. Colonies were picked up after 5, 10 and 20 days during the reprogramming process (Figure 1a).

**Figure 1.**
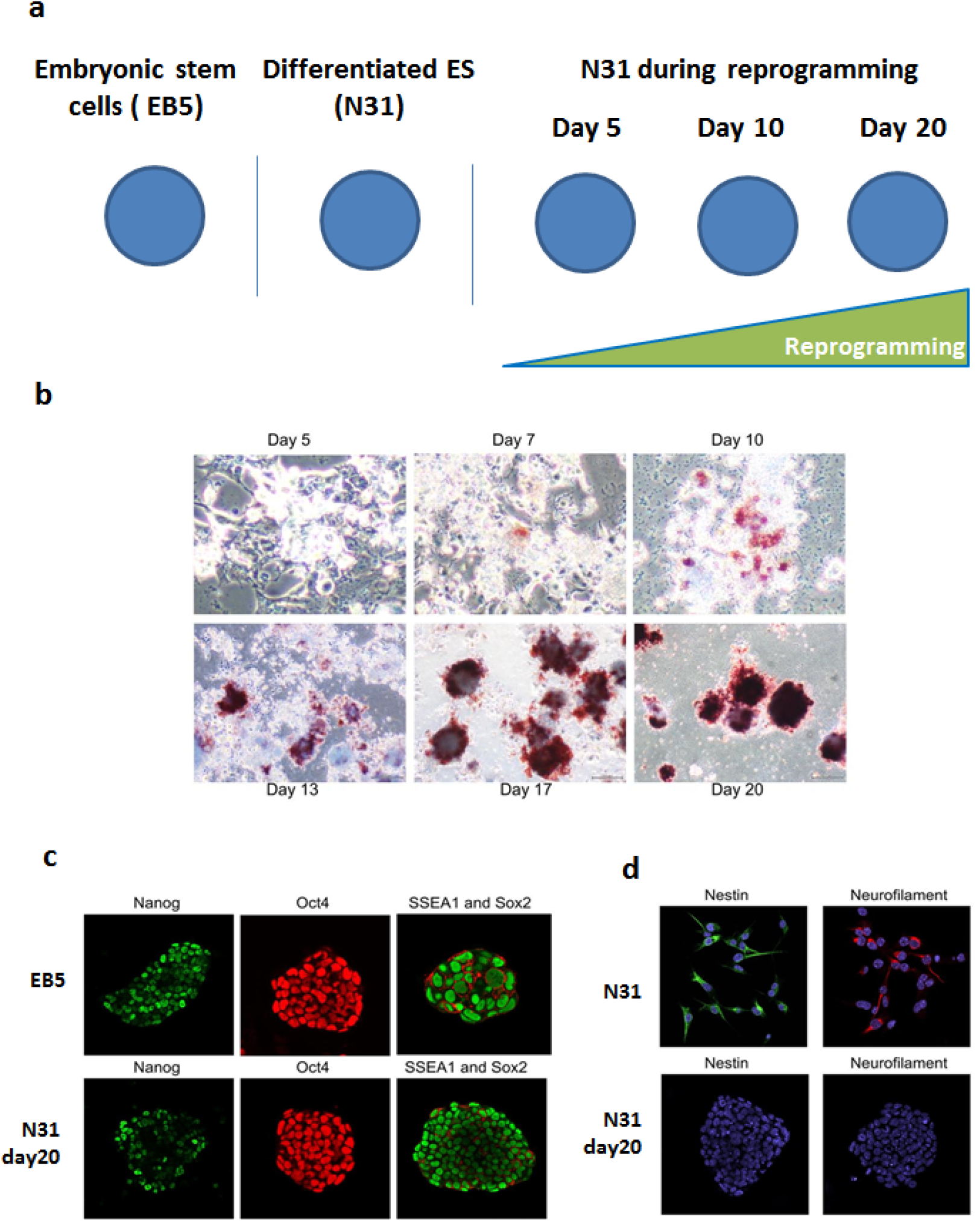
Cell states derived from the original stem cells, and assessment of reprogramming by AP staining and immunostaining. **a)** ES stem cells EB5 and its neural progenitors N31 make an isogenic cell line. N31 was then subjected to reprogramming for 20 days, and Raman spectral measurements were performed at three time points during reprogramming. **b)** Alkaline phosphatase staining of N31 cells during the reprogramming process. Dark red indicates positive. staining. **c)** Immunofluorescence analysis of EB5, and N31 cells after 20 days of reprogramming. Both exhibit colonies which are positive for pluripotent markers Nanog, Oct4, Sox2 and SSEA1. **d)** Immunofluorescence analysis for neural progenitor markers Nestin and Neurofilaments of the differentiated cells N31and reprogrammed iPS-like colony obtained 20 days (upper row - N31, lower row - 20 days after reprogramming). Nuclei stained with DAPI appear in blue.

### Immunostaining

For immunofluorescence analyses cells were grown on glass bottom 30-mm dishes coated with collagen type I (IWAKI #4970-011). On the day of immunostaining cells were briefly washed with PBS, fixed with 4% PFA (Santa Cruz #sc-281692) for 15 min at room temperature and permeabilized with 0.5% Triton in PBS with 10% FBS addition for 30 min. Primary antibodies were applied: Anti-Oct 3/4 (Santa Cruz #sc-5279, 1/250 dilution), Anti-Nanog (Abcam #ab80892, 1/250 dilution), Nestin, Neurofilament in PBS with 10% FBS addition, for 1 hour in room temperature. After washing cells were incubated with secondary antibodies: Anti-mouse Alexa Fluor^®^ 594 (Cell Signaling #8890, 1/500 dilution) and Anti-rabbit Alexa Fluor^®^ 488 (Cell Signaling #4412, 1/500 dilution) in PBS with 10% FBS addition for 1 hour at room temperature, then cells were washed 4 times with PBS and 2ml PBS per dish was added for imaging.

### Alkaline phosphatase staining and imaging

For alkaline phosphatase staining cells were briefly washed with PBS, fixed for 5 min with 4% PFA (Santa Cruz #sc-281692) at room temperature and stained with Alkaline phosphatase kit II (Stemgent, #00-0055) according to the manufacturer’s protocol. Imaging was carried out on Olympus CKX41 inverted microscope (Olympus, Tokyo, Japan).

### Raman spectroscopy and spectral pre-processing

A homemade confocal line-scanning Raman microscope was used, which optical system is described in details in a previous study ^7^. The spatial resolution of our system is approximately 300 nm, and spectral resolution approximately 1 cm^-1^. A 532 nm diode-pumped solid-state laser (Ventus, Laser Quantum, UK) was focused through a water-immersion objective lens (NA: 1.20, UPLSAPO60×W, Olympus, Tokyo, Japan). In this study, cells were kept alive, which contrasts with several previous studies ^15,16^. Glass bottom dishes were placed onto a heated micro-chamber, set at 37°C and 5% CO_2_ to encourage the cell survivability and avoid metabolic stress. Single living cells were line-scanned using a 5 sec laser exposure per line and only 10 lines per cell were taken, so as to accelerate measurements. Measurements were performed using WinSpec (Princeton, New Jersey, USA) synchronized to a home-made automated program developed in IGOR (IGOR Pro v6, WaveMetrics, Inc., Portland, US). Raman spectral data were first treated to remove cosmic rays. For each single cell image, the pixels corresponding to the cell were averaged, and the pixels corresponding to background were averaged. After background subtraction, a single spectrum is obtained for each cell. Then, a polynomial baseline correction using the ModPoly algorithm ^17^ was applied on background subtracted spectra for the range 550 to 1750 cm^-1^. Because Raman spectra measured on different days can produce slight shifts, due to the imperfect reproducibility of the grating angle in the polychromator, data were interpolated using a cubic spline function allowing to recalculate the x-axis. Then edges were cut-off, to give spectra for the range 590 to 1710 cm^-1^ with 1129 variables. Then the spectral data were vector normalized. These pre-processing steps were conducted using homemade functions in MatLab (MatLab 2015a, Mathworks, US). Local maxima of specific Raman bands were identified for each single cell by using a home-made algorithm using a pre-given range of search around known wavelengths.

### Multivariate analyses

#### Neural Network classification model

To determine if the spectral variations were significant among strains, classification analysis was performed using a Neural Network model. classification was performed using 1 dimensional convolutional Neural Network (1DCNN) which is composed of convolution layers, max pooling layers and fully connected layers (Figure S1). Batch normalization ^18^ and dropout ^19^ were also used to enhance the robustness of the prediction results. In the training and prediction process we adopted 8-fold nested cross validation, in which all data was divided into 8 datasets and the ratio of training data, verification data, and test data was distributed at 6: 1: 1. There were a total of 56 ways to divide by the distribution ratio, and 56 types of 1DCNN models were trained. In this case, since 7 models were trained on 1 test dataset, and 7 predictions were obtained for each data. Through the above processes, prediction of the 1DCNN was performed for all data. To visualize the wavenumbers that contributed to the classification of each cell state, a score was extracted. After training with all the data and the same model as 1DCNN used for the prediction, the wavenumber was output to all the data by Grad-CAM ^20^.

#### PLS-DA models

While the Neural Network model aimed at classifying the five cell states from each other, we then wanted to visualize the wavenumbers contributing to different cell states “direction”. To do so, we computed two distinct low dimensional spaces using Projection on Latent Structure-Discriminant Analysis (PLS-DA) model ^21^, by considering pairs of two cell states: EB5/N31 in the first model, and N31/N31d20 in the second, respectively. In this the twodimensional space, the F1 axis represents the direction that best separate the two types of cells. The Variable Importance on Projection (VIP) scores, which represents the importance of each wavenumber in separating the cell types, were also calculated. Prior to analysis, normalized spectral data were mean centered. A 100-splits venetian blind cross-correlation method was applied. The model complexity (the number of components in the model) was determined by using the complexity that minimized the RMSECV value. Model complexity was found to be 4, for the comparison EB5/N31, and 2 for the comparison N31/N31d20. Then, with the hypothesis that ES cells and differentiated cells define “two ends” of the range of possible cell-state, we wanted to see how the intermediary states would be defined along this axis. We used the above PLS-DA trained from the Raman spectral data of the ES (EB5) and differentiated cells (N31). In this low dimensional space, we then projected the spectrum of cells during reprogramming (N31d5, N31d10, N31d20) in the already built space (as test data) and compared the distribution of F1 scores of all cell states. Models were computed using the PLS toolbox in Matlab (EigenVector, Manson, USA).

## Results

### Confirmation of cell states by AP staining and immunofluorescence

The Figure 1a describes the experimental layout and the cell states induced and assessed in a mouse isogenic cell line called EB5. As shown in the figure, EB5 cells were differentiated in vitro to create neural-lineage N31 cells, and then N31 cells were iPS-reprogrammed, and samples were taken throughout the process on days 5, 10 and 20 for Raman imaging. Thus, all cells used in this study are isogenic progeny of the original ES cell line EB5 and can be referred to as “cell states”. Figure 1bcd shows the alkaline phosphatase (AP) and immunofluorescence analyses of these cell states, with immunofluorescence and AP staining serving as conventional methods to verify the pluripotency status ^22^. Figure 1b illustrates the gradual changes in AP positivity in N31 cells during the reprogramming process, confirming that N31 cells underwent the reprogramming as intended. Figure 1c shows the immunofluorescence analyses of the original ES cell line EB5 and of N31 cells after 20 days of reprogramming (N31d20), confirming that both ES and N31d20 colonies were positive for pluripotent markers Nanog, Oct4, Sox2 and SSEA1. Figure 1d shows the immunofluorescence analysis of differentiated cell line N31 demonstrating the presence of neural progenitor markers characteristic of neural lineage cells ^22^ and absence of these markers from N31d20, confirming that N31 cell line was differentiated, while N31d20 lost these differentiation markers after 20 days of iPS reprogramming.

### Label-free Raman spectroscopy highlights cell-state-specific signatures

The state of living single was then evaluated by Raman spectroscopy. About 40 peaks were clearly visible from the spectral signature. The molecular peak assignment of spectral data was performed according to previous studies ^7, 11, 13^. In Figure 2a, the averaged normalized spectra were compared for visualization purposes. These data highlighted strong differences between the cell states derived from ES cells, notably, the peaks at ~718 cm^-1^ (CH, CS quinolo), ~752 cm^-1^ (Tyr, CC, cytochrome), ~786 cm^-1^ (PO_2_ stretch of DNA, cytosine), ~ 989 cm^-1^ (ß-sheets of protein), ~1260 cm^-1^ (nucleic acids, Amide III of lipids), and ~1445 cm^-1^ (CH_2_ deformation - lipids and proteins), highlighted strong differences between ES cells and cells undergoing reprogramming and their progenitor N31. Despite variations of spectral intensities within each cell state, the differences of the local maxima of each band at the aforementioned wavelengths were found significant (*p* < 0.05, ANOVA followed by a post-hoc Tukey HSD). In particular, stem cells exhibited high intensity for fatty acids (double bond stretching at ~1260 cm^-1^ and ~1650 cm^-1^) and lower amounts in unsaturated lipids (~1445 cm^-1^) than other cells. The differences between ES cells and N31d20 are highlighted in Figure S2.

**Figure 2.**
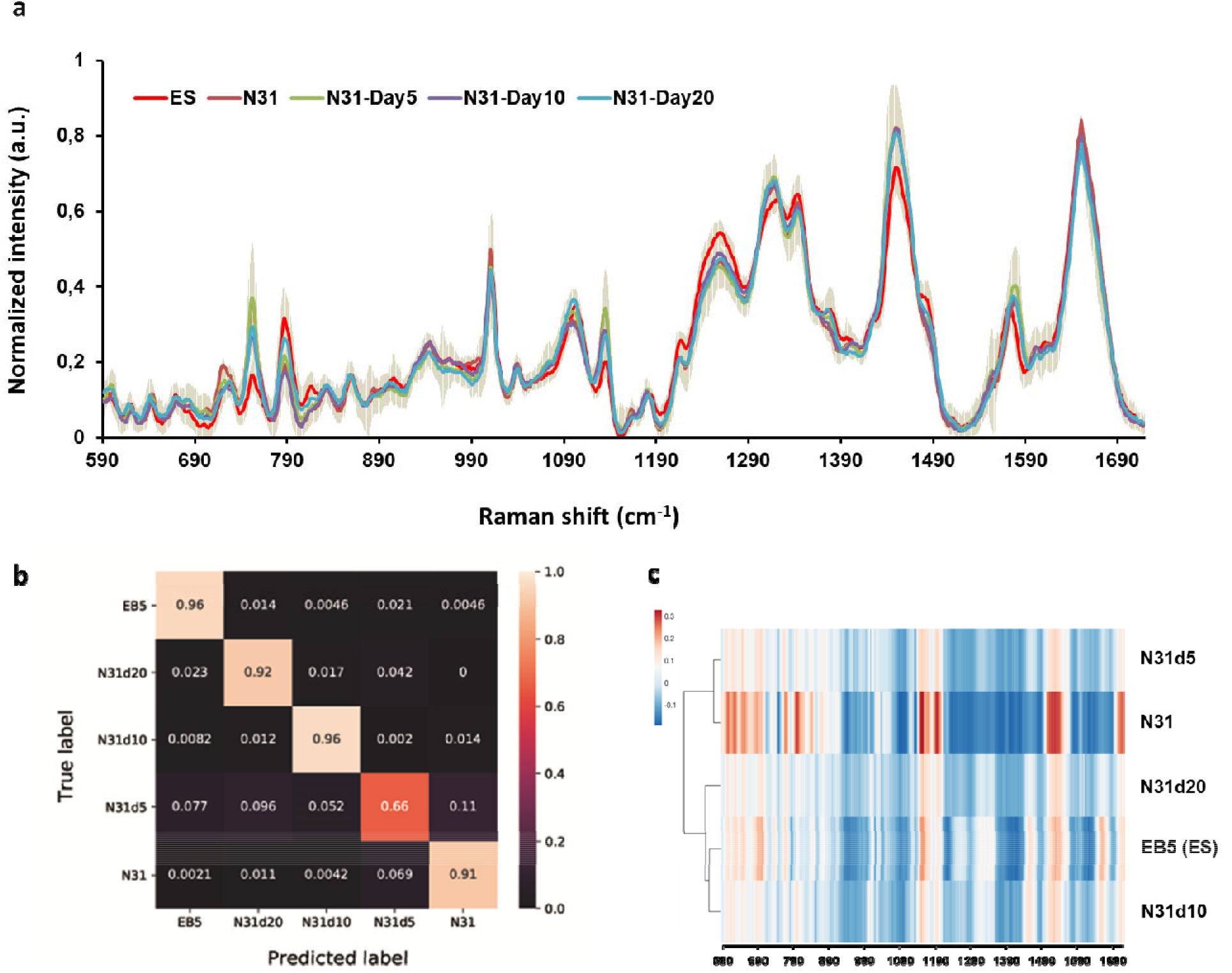
Comparison and prediction of cell states from single cell spectral data by machine learning. **a)** Normalized averaged spectral data of single cells across the five cell states. ES cells (EB5 n = 62), neural cells (N31 n = 68), and cells undergoing differentiation for 5 days (N31d5, n = 61), 10 days (N31d10 n = 71) and 20 days (N31d20 = 68). **b)** Confusion matrix for the predictions of 1DCNN versus the true labels. The numbers represent the average probabilities of the 7 times predictions due to 8-fold nested cross validation in each class. **c)** Hierarchical clustering (Euclidian, average) of the weights associated to each wavelength extracted from the neural-network classification. The result shows the similarities between N31 and N31d5, and similarities between ES cells EB5 and N31d10 and N31d20.

A neural network-based classification model was developed to measure the ability to predict each cell state based on the whole Raman spectra of single living cells (Figure 2b). The dataset was divided into training and test dataset by holding out 20% of the spectra. Each cell state was discriminated against each other with a very good accuracy, ranging from 0.66 to 0.96%, and an average accuracy of 88.2%. This result demonstrate that Raman spectroscopy data are very rich to allow the prediction of the cell state by CNN. The lowest score of 0.66% was observed for N31d5 data, which highlight a stronger variability in the molecular signatures of these cells. To visualize the similarities between cell states, the weights associated to each wavelength using the output of Grad-CAM (Supplementary Data) were used to compute a hierarchical clustering (Figure 2c). Figure 2c shows the N31 and N31d5 were clustered together, suggesting their close similarities in spectral profiles. ES cells (EB5) were clustered with cells after 10 or 20 days of reprogramming (N31d10, N31d20) suggesting similarities. This clustering results valid the reprogramming process is effective and push N31d10, N31d20 cells towards the EB5 stem cell state.

### Exploring possible spectral biomarkers of differentiation and reprogramming

To highlight the spectral bands that can potentially account for differences during the process of differentiation or reprogramming is of interest to this study. We therefore performed a data-driven analysis of ES cells (EB5) against ES-derived differentiated cells (N31) by computing a PLS-DA, a classification model, using all samples from these two cell states (Figure 3a). The spectral data were projected in a low dimensional space built on two-components. Our analysis shows the cross-validated model can identify these two cell states from their F1 scores (Figure 3a, left panel). The F1 vector (Figure 3a, right panel) highlights the wavelengths of importance to classify into one cell state or the other. We also extracted the VIP scores which calculates the contribution of each wavenumber in identifying these strains (Figure S4). Both F1 score and VIP scores matched the local maximum of known Raman bands, allowing us to identify the peaks representative of the cell states. Notably, the bands at ~752 cm^-1^ (Tyr, CC, cytochrome), ~1002 cm^1^ (phenylalanine - protein), ~1133 cm^1^ (C-H in-plane bending mode of phenylalanine, cytochrome proteins), ~1260 cm^1^ (nucleic acids, Amide III of lipids) and ~1445 cm^1^ (CH_2_ deformation - lipids and proteins) strongly contributed to identify neuronal cells (N31). Oppositely, the band at ~786 cm^-1^ and ~1240 cm^-1^ were important to characterize ES cells.

**Figure 3.**
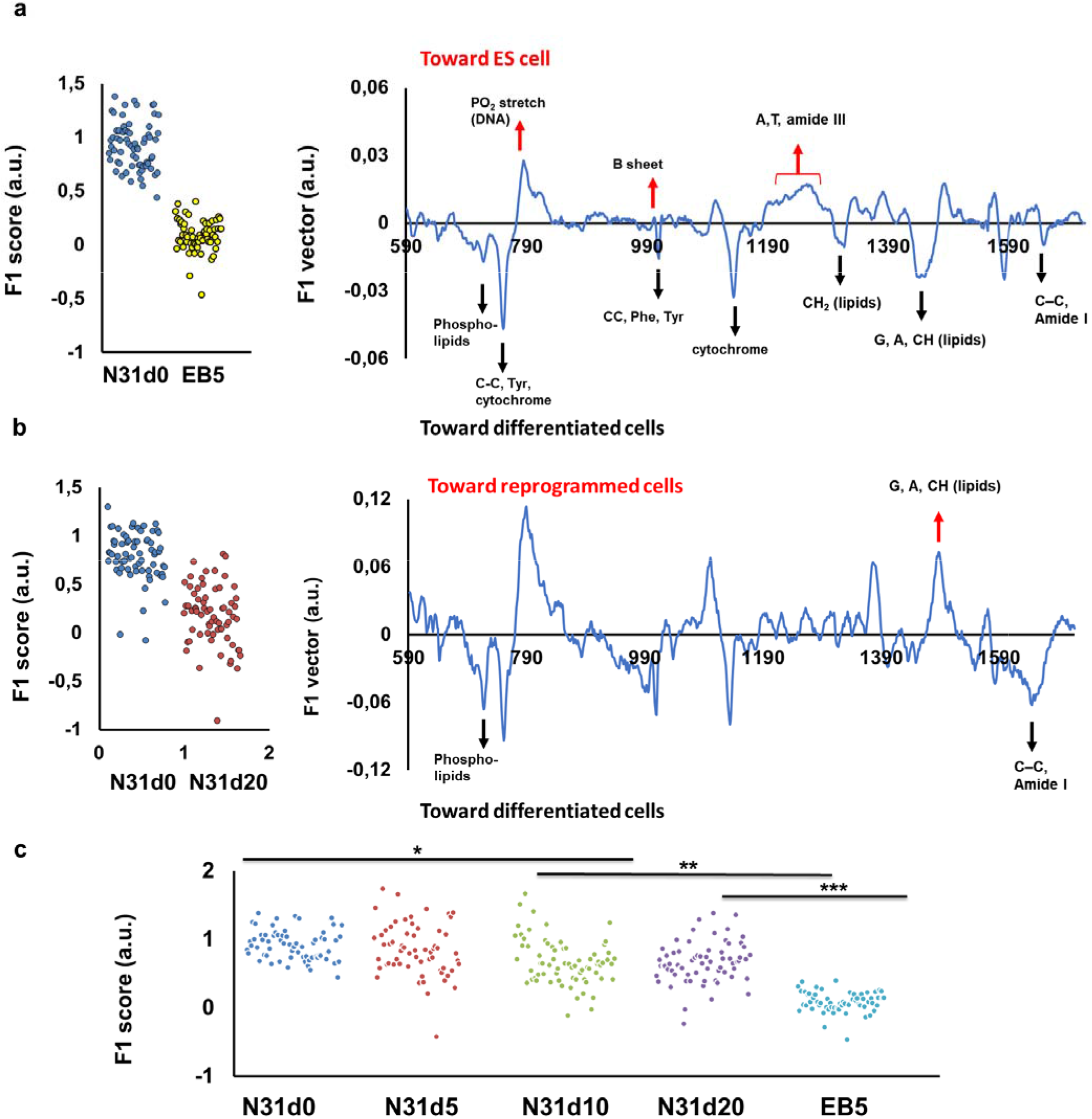
Cell-states and characteristic analysis by PLS-DA. **a**) PLS-DA trained using ES cells (EB5, n=62) and its neural progenitor (N31, n= 68). The F1 scores (left panel) extracted from the model show the possibility to identify the two cell-states along the F1 vector (right panel) which show the characteristic wavelengths of importance to identify each cell state. A few important peaks are shown. Horizontal axis of right panel are expressed in Raman shifts (cm^-1^). **b**) A PLS-DA model was trained using N31 cells (n=68) and its reprogrammed counterpart (N31d20, n= 68). The F1 scores (left panel) and F1 vectors (right panel) specific to this model are shown. X axis of the right panel are Raman shifts (cm^-1^). **c**) Using the low dimensional space built in Figure 3a, spectral data of cells undergoing reprogramming (N31d5, N31d10, N31d20) were used as test data and plotted in the already-built space. The F1 scores extracted from the model reveal the possibility to identify each cell state along the continuum N31-EB5. Asterisks over groups show statistical differences between indicated pairs, according to an ANOVA followed by a post-hoc Tukey HSD test (*p* < 0.05).

Similar to the above, we compared the differences between N31 and N31d20 by computing a PLS-DA model with these two cell states (Figure 3b) to search for potential spectral biomarkers accounting for the reprogramming process of neuronal cells toward IPS-cell state. The F1 vector shows the peaks contributing to identify neuronal cells (N31) are similar than in the above model, at the exception of the 1445 cm^-1^ (CH_2_ deformation - lipids and proteins) which this time strongly contributed to identify reprogrammed cells (N31d20).

With the possibility that N31 and EB5 are two “ends” of the cell state transition in isogenic cell state, we verified how spectral data of cells undergoing reprogramming (N31d5, N31d10, N31d20) would position along this axis. We then used the data of cells undergoing reprogramming (N31d5, N31d10, N31d20) as test data which were plotted into the already-built dimensional space defined by stem cells and neuronal cells (Figure 3a). We found that the F1 scores of the data classify these cells state at intermediary scores. The distribution of the F1 scores for each cell state follows a normal distribution (data not shown). The F1 scores N31d10, N31d20 and EB5 were statistically different from the other cell states.

### Ratiometric analysis

The above machine learning-based analyses reveal spectral data contain invaluable features to characterize the differentiation and reprogramming process. To describe differences in cell states, one can also calculate the ratios between the intensity of the peaks identified in the above analyses. Table 1 gives a list of a few ratio values we explored, which also included ratios described in previous literature. Ratio were calculated using the local maxima of peak of normalized spectrum, for each sample. Ratios were arbitrarily annotated from R1 to R8. Ratiometric data and results of statistical tests performed using single cell data, are provided in Supplementary Data. Ratios were plotted in Figure S5 and S6 with their standard deviation. Several of the shown combinations were particularly efficient at discriminating cell states, at single-cell level, with significant *p* values. In particular, the average value ratio for ~752 / ~786 cm^-1^ provided the strongest change across cell state, specifically a 4-time change, as the average value ranged from 2.02 for N31 to 0.56 for EB5, with intermediary values for cells undergoing reprogramming. A post-hoc Tukey HSD test highlighted significant (p < 0.05) differences between the states N31d5, N31d10, N31d20 and EB5. However, no statistical difference was found between N31 and N31d5 using this specific ratio. In addition, the ratios R1 (~718 / ~756 cm^-1^) and R5 (~1260 / ~1445 cm^-1^) allowed to distinguish EB5 against the others (Supplementary Data) but did not allowed to distinguish N31 against the reprogrammed cells. For several ratio pairs, the N31 and EB5 showed the strongest or weakest values by comparison to cells undergoing reprogramming (Supplementary Data).

**Table 1.**
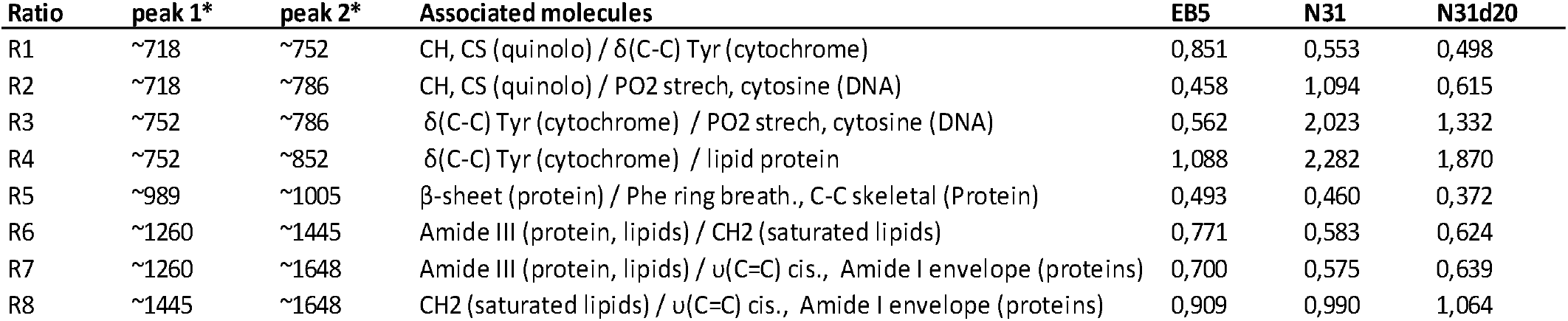
Ratiometric analysis of bands of interest during reprogramming and differentiation in mouse cells. Spectral intensity values were obtained from the local maximum. *Peak values are expressed in Raman shifts (cm^-1^). For each ratio, at least one cell state was statistically different from the others (ANOVA, post-hoc Tukey HSD p < 0.05). Standard deviations and statistical groups identified by a post-hoc Tukey HSD test are not shown in the table for clarity but are provided elsewhere (Supplementary Data, Figure S5, S6).

## Discussion

It is well known that stem cells undergoing differentiation undergo deep changes in their metabolic properties and gene expression, and to monitor such processes in a non-invasive manner, novel bioimaging techniques are particularly desirable. Using a modest single-cell dataset of mouse ES, differentiated neuronal cells, and cells undergoing reprogramming, this study exemplified that Raman spectral characteristics and machine learning allow to monitor cell-state transition in an isogenic cell line, while providing biologically-relevant biomarkers to characterize these cells states. Our classification model based on a neural network shows a good accuracy in identifying unknown (new) cells from their chemical profiles (Figure 2bc). This shows the ability of Raman spectroscopy to discriminate accurately each cell state, and even predict the states from the spectral measurement of unknown cells using a trained model. A hierarchical clustering performed using the output of Grad-CAM (Figure 2c) seemed to reflect the biological proximity of cell-states, such as N31 and N31d5 which clustered together and were significantly different from the other states. Interestingly, a similar result was also obtained by building a low dimensional space with ES and neuronal cells and plotting other cell states in this space and extracting their F1 scores (Figure 3c). Therefore, one key result of this study is that the chemical or metabolic properties of cells captured by Raman spectroscopy could define a low dimensional space in which the different cell states were visualized, monitored and characterized. While the ability to monitor the dynamic of differentiation in a different cell line has been reported previously ^10–13^, a study of cell states during the reprogramming process using an isogenic cell line is reported for the first time.

Potentially, prediction scores or F1 value distribution can also highlight which cell-state is the most difficult to characterize. N31d5 cells, in particular, had a strong lower score in the CNN model (Figure 2b) and strong variation in F1 scores (Figure 3c). Hypothetically, this low score may be due to the inherent large heterogeneity of metabolic state in the early reprogramming process, with the expectation that while some cells started the reprogramming, others didn’t. A similar result was found by Ichimura and colleagues ^10^ for an “intermediary” state during differentiation. To further investigate whether Raman spectroscopy is sensitive enough to monitor the cell state transition in early reprogramming, additional experiments should be carried out by measuring cells every day or every 2 days.

This study also identified the so-called spectral biomarkers using several methods such as spectral differences (Figure S2), F1 vector analysis (Figure 3ab, Figure S3) and ratiometric analysis (Table 1, Figure S5). Ratio of spectral intensities are powerful tools when analyzing Raman spectral data in the sense that they provide an “unbiased” way of exploring the data. In fact, one can expect that ratio values to be conserved across different platforms with different optical setups (despite possible differences in background level or sensitivity), if at least the excitation wavelength remains the same. In previous studies, Schulze and colleagues proposed a ratio of the differentiation state marker, specially using the ~752 cm^-1^ and ~786 cm^-1^ bands in human cells ^13^. However, in this pioneer work, only a few cells were used, and the spectral area that was investigated was limited to 663-1220 cm^-1^. In our study, all ratios shown in Table 1 could significantly distinguish between specific pairs of cell states. The ratios R1 (~718 / ~756 cm^-1^) and R5 (~1260 / ~1445 cm^-1^) for instance, allowed to distinguish EB5 against the others (Supplementary Data). The ratio R3 (~752 /~786 cm^-1^), associated with cytochrome/proteins, was extremely valuable to monitor both the differentiation and the reprogramming process and can be considered as a good spectral biomarker.

The ratio between ~1260 / ~1445 cm^-1^ (Amide III and Amide I, respectively) highlight changes in unsaturated/saturated lipids, as the first band is an important indicator for unsaturated acids (such as DAG), which get stronger during the loss of unsaturation such as oxidation, and the second an important marker for saturated fatty acids ^23^. By comparison to neuronal cells, the lower ratio for EB5 cells suggests it has more unsaturated fatty acids than saturated ones. The Figure 1a also suggests it as EB5 clearly displays a lower intensity value at ~1445 cm^-1^ by comparison to other cell states. Interestingly, this observation is consistent with the fact that ES cells have high amount of ω-6 and ω-3 polyunsaturated fatty acids (e.g., arachidonic acid, docosahexaenoic acid and linoleic acid) which are quasi-absent in differentiated cells and low in cells undergoing reprogramming ^24^. The spectral variations at ~1260 and ~1445 cm^-1^ seem therefore useful to monitor the reprogramming and differentiation process. Further work must be conducted to show how lipids bands correlate with fatty acid synthesis, which is known to be critical for stem cell pluripotency^25^.

Interestingly, the above results hint at differences between ES and cells undergoing reprogramming (N31d20), even at later stages (N31d20, positive for pluripotency markers Nanog, Oct4, Sox2 and SSEA-1). We cannot fully exclude the possibility that the differences between ES cells and N31d20 are due to the incomplete iPS reprogramming or the heterogeneity of N31d20 cells. We should highlight, though, that the immunofluorescence analysis showed no differences in expression of pluripotency markers Nanog, Oct4 and Sox2 in ES and N31d20 cells. Nevertheless, the spectral differences at 1445 cm^-1^ (Figure S2, Figure 1a) make a strong case for substantial differences between these cell states. Again, fatty acid synthesis is known to be critical for stem cell pluripotency^25^. Thus, the observed spectral difference is of interest with regards to medical applications of iPS cells such as transplantation ^2,3,4^, as the quality or safety of iPSCs is often performed by comparing the cells to naturally occurring embryonic stem cells (ESCs). Yet, whether the iPS cell state constitutes an equivalent to naturally occurring ES pluripotency is an undergoing debate in the community. Numerous attempts have been made to find differences that distinguish ESCs from iPSCs, identifying dissimilarities in gene expression patterns ^26–30^ DNA methylation ^31–33^) and chromatin modifications ^30, 34^. Comparisons with label-free imaging data are therefore desirable and in the light of our results, we speculate that spectral methods have a strong potential in highlighting differences between ES and reprogrammed cells.

To conclude, this study showed the possibility to discriminate the differentiation and reprogramming process of living cells by label-free spectral measurements, showing the richness and unique information brought by vibrational spectroscopy. Our data reports for the first time several important biomarkers specific to differentiation or reprogramming in mouse cells, which confirm or complement previous studies on human investigations. This gives a strong basis for selecting the wavelengths identified in this study to develop high-throughput label-free identification pluripotency or cell-state dynamics, by selecting a pre-given number of wavelengths or ratios. Additionally, we demonstrated how cells undergoing reprogramming exhibited different states in a low dimensional space built upon ES and neuronal cells signatures, and it could be interesting to evaluate how the cell-state defined by different modalities can complement each other’s. As a perspective, we expect that data-integration of Raman data (image, or spectra) can be done with complementary modalities, such as metabolic analyses (eg. ^27^), and that will extend our understanding of the biological causes for spectral shifts observed between cell-states. We hope this modest study will become a valuable reference for future studies of cell-state investigations in differentiated and iPS cells.

## Link to datasets

Our datasets and analysis reports are provided into an excel file for convenience, available at XXXXX.

## Author Contribution

A. G. did the experiments, data analysis, and wrote the paper. Y. P. did experiments and writing. M.S. and H.N contributed to multivariate analysis. T. M. W. provided critical comments.

