## Supplementary information for "Following embryonic stem cells, their differentiated progeny, and cell-state changes during iPS reprogramming by Raman spectroscopy"

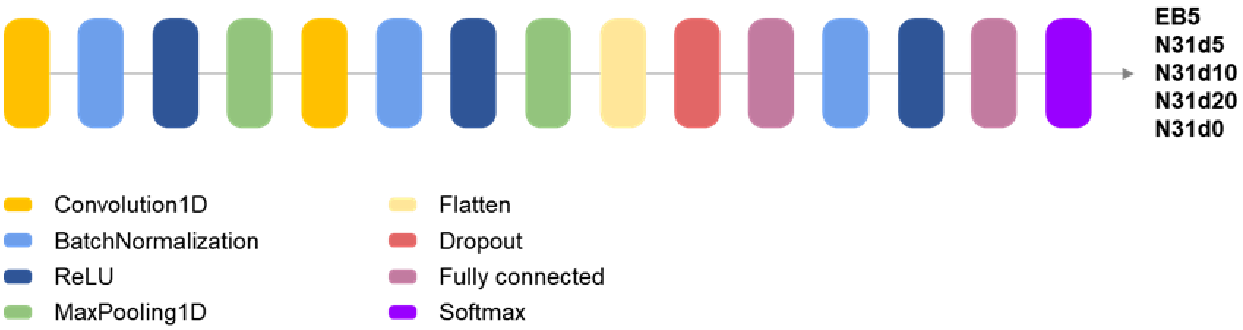


**Figure S1. Structure of the neural network model.**

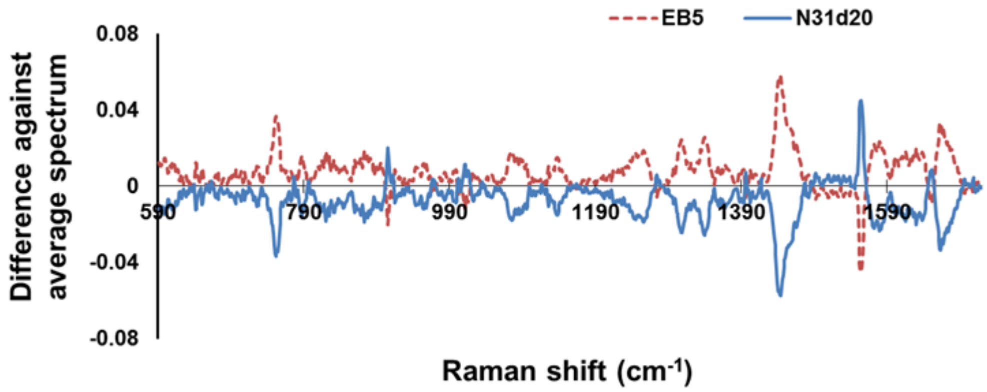


**Figure S2. Differences of EB5 and N31d20 spectra against their average spectrum.**

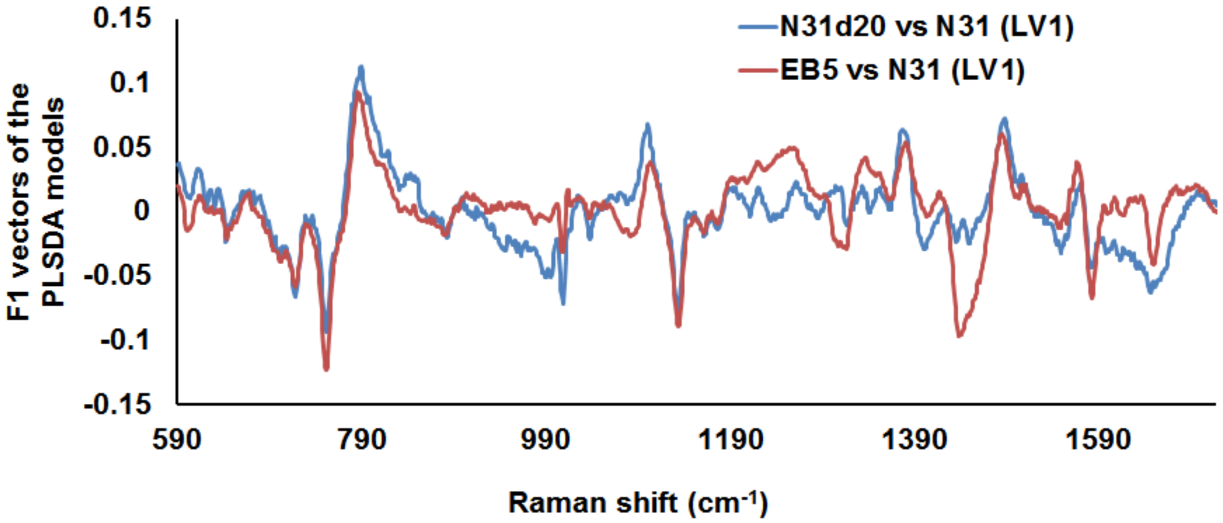


**Figure S3. Overlapped F1 vectors of the two PLS-DA models.** This visualization reveals a particularly strong difference in the contribution of the 1445 $cm^{-1}$ Raman band in each model.

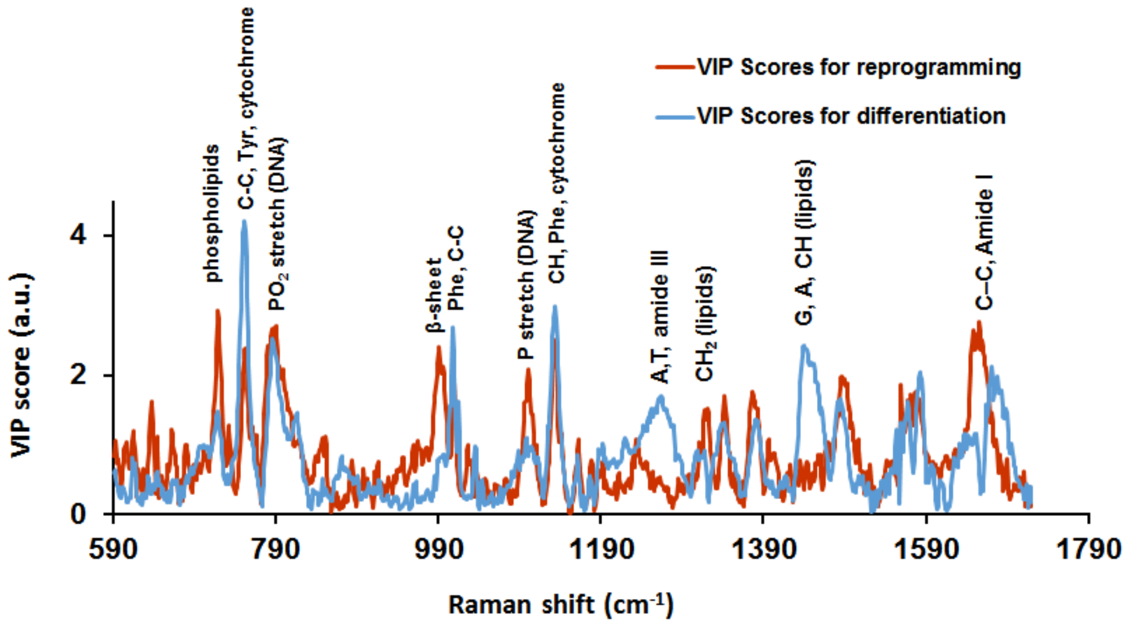


**Figure S4. VIP scores associated with the wavenumbers (Raman shifts) which contributed the most in classifying the cell state in PLS-DA models.** Model for reprogramming was trained using ES cells (EB5, n=62) and its neural progenitor (N31, n= 68), and the model for reprogramming was trained using N31 cells (n=68) and its reprogrammed counterpart (N31d20, n= 67). Only major peaks with VIP scores values above 1 are considered significant.

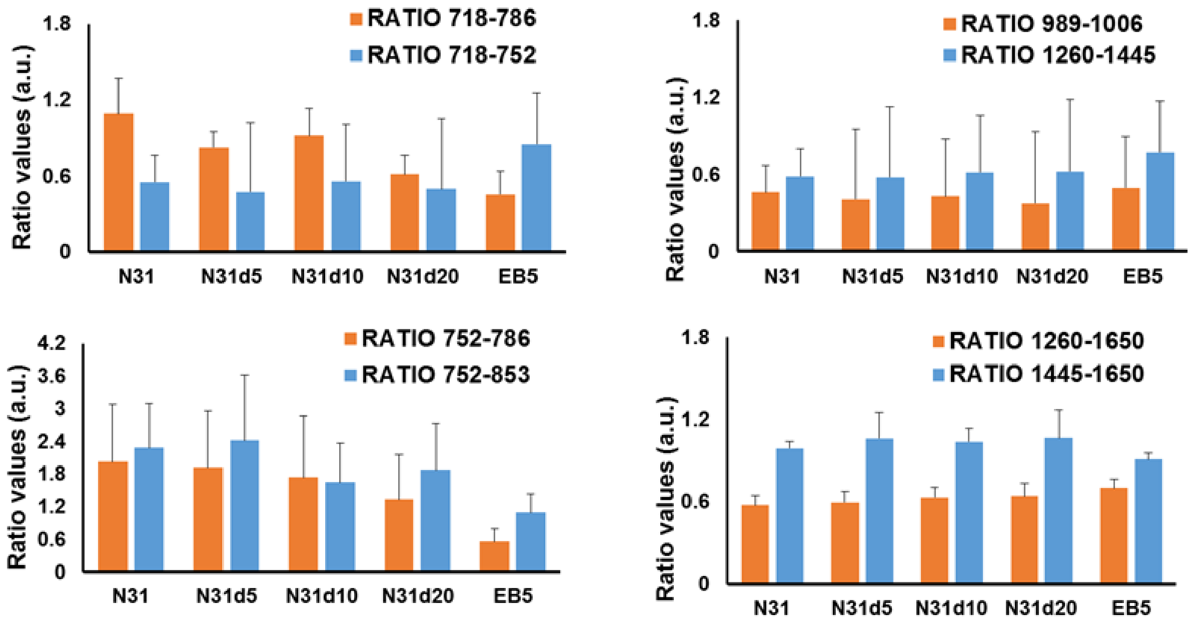

**Figure S5. Histogram visualization of the ratiometric analysis for the 8 pairs of ratio described in Table 2.** Histograms are the average and vertical bars the standard deviation (SD) calculated across all single-cells for each cell-state.

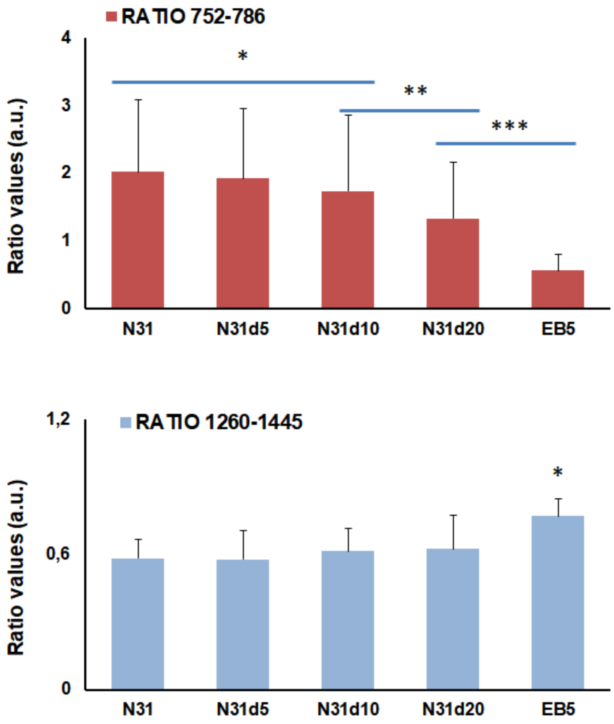


**Figure S6. Histogram visualization of the ratiometric analysis for two pairs of ratios described in Table 2.** Histograms are the average and vertical bars the standard deviation (SD) calculated across all single-cells for each cell-state. Asterisks represent statistical differences according to ANOVA followed by post-hoc Tukey HSD test (p 0.05).
